# Exposure to fruit-flavoring during adolescence increases nicotine consumption and promotes dose escalation

**DOI:** 10.1101/2021.01.12.426210

**Authors:** Theresa Patten, Allison Dreier, Rae J. Herman, Bruce A. Kimball, Mariella De Biasi

## Abstract

The rise of e-cigarette popularity has sparked interest in the role of palatable flavors on nicotine use. Despite growing evidence that sweet flavorants enhance nicotine reward, their influence on nicotine consumption has not been studied extensively. In addition, the impact that flavored nicotine use in adolescence could have on nicotine reward and dependence in adulthood remains unclear. This study examined the role of flavored nicotine access on nicotine preference and consumption longitudinally, from adolescence to adulthood. Male and female adolescent mice preferred a fruit-flavored nicotine solution. However, only adolescent female mice with access to flavored nicotine consumed higher doses. Furthermore, while adolescent male mice escalated consumption of both flavored and unflavored nicotine, female mice only escalated when given access to flavored nicotine. As mice matured into adulthood, there was no evidence that a history of flavored-nicotine access altered preference for unflavored nicotine. However, when the nicotine concentration was reduced, mice that had consumed strawberry-flavored nicotine in adolescence maintained baseline nicotine consumption levels longer than mice that initiated nicotine use without flavor in adolescence. Finally, addition of fruit-flavorants into the nicotine solution during adulthood led to increased levels of nicotine consumption, regardless of previous flavored-nicotine access or of familiarity with the selected flavorant. These results indicate that flavorants increase nicotine consumption independent of life stage, possibly posing a disproportionate risk to adolescent females. Our results also point to an effect of adolescent flavored-nicotine use on nicotine dose maintenance in adulthood, which could have implications for the success of future quit attempts.

## 1. Introduction

E-cigarettes arrived on the market in the early 2000s and they have been increasing in popularity among adolescents for the past decade (Gentzke et al., 2019; National Academies of Sciences, et al., 2018). E-cigarettes are available in over 15,000 distinct flavors that are attractive to adolescents (e.g. ‘cotton candy’, ‘tropical blue slushie’ and ‘crazy berry’) (Hsu et al., 2018). These sweet and ‘characterizing’ flavors have played a major role in e-cigarette uptake and popularity among young people, with nationally representative data sets regularly showing that ‘flavor availability’ is among the top two reasons for e-cigarette use and experimentation in students (Ambrose et al., 2015; Bold et al., 2016; Kong et al., 2015; Patten and De Biasi, 2020).

The flavoring compounds used by e-liquid manufacturers are the same as those that children grow up consuming in candy and sugary drinks (Brown et al., 2014). As a result of repeated pairing with highly palatable and caloric foods, these flavorants have established positive associations well in advance of nicotine exposure (Beauchamp and Cowart, 1985; Fanselow and Birk, 1982; Harris et al., 2004; Mennella et al., 2016). Compared to adults, adolescents have a heightened preference for both sweetness and for the foods and flavors paired with sweetness (Cooke and Wardle, 2005; Desor and Beauchamp, 1987; Hoffman et al., 2016; Mennella et al., 2016). Therefore, it is not surprising that sweet and fruit-flavored e-cigarettes are overwhelmingly attractive to adolescents. In fact, between 80-98% of adolescents and 60-95% of young adults initiate e-cigarette use with a flavored e-cigarette, underscoring flavor’s importance in e-cigarette use among these populations (Harrell et al., 2017; Villanti et al., 2017). In this context it is important to note that the 2020 FDA ban on flavored e-liquid cartridges (https://www.fda.gov/) only affects cartridge-based products such as JUUL while the sales of flavored e-liquids designed for open-tank systems and disposable cartridge-based products remain unaffected.

In addition to their role in e-cigarette uptake, flavorants may impact nicotine reward and consumption during e-cigarette use. Characterizing flavors enhance subjective reward of nicotine-containing e-cigarettes and can increase vaping patterns, such as number of puffs taken, volume of e-liquid used, and duration of puffs (Audrain-McGovern et al., 2016; Goldenson et al., 2016; Jackson et al., 2020; Kim et al., 2016; Leventhal et al., 2019a; St.Helen et al., 2018). However, changes in these vaping behaviors do not easily extrapolate to total nicotine exposure. For example, in one study, although longer puff durations were observed when participants vaped a strawberry-flavored e-cigarette, increased puff duration was not associated with a statistically significant increase in systemic nicotine exposure (St.Helen et al., 2018). Another study in adult male rats showed that licorice flavor enhances nicotine self-administration (i.e. consumption). However, this study did not address how flavors (and more specifically, how flavors popular with young people) could impact nicotine intake during adolescence, a period when users are particularly vulnerable to both nicotine and flavorings (Palmatier et al., 2019). Notably, the study by Palmatier and colleagues (Palmatier et al., 2019) included only male subjects. Women show a slightly higher preference for- and use of-e-cigarettes with characterizing flavors, and they may be more sensitive to the sensory components of vaping/smoking (Kistler et al., 2017; Patten and De Biasi, 2020; Perkins, 1999; Perkins et al., 2001; Soneji et al., 2019; Xiao et al., 2019). Overall, there is a need to better understand the influence of flavored additives on nicotine intake levels, especially in females and in adolescents.

Individuals who report a positive first-experience with smoking are more likely to go on to become regular smokers (Chen et al., 2003; DiFranza et al., 2007; Mantey et al., 2017; Rodriguez and Audrain-McGovern, 2004; Sartor et al., 2010; Urbán, 2010). Additional research suggests that reducing the initial aversion to a bitter taste (such as that of nicotine), could reduce aversion to nicotine if and when adolescents transition to combustible tobacco use (Capaldi and Privitera, 2008; Hoffman et al., 2016). It is a public health concern that this generation of young ‘vapers’ could experience more severe long-term consequences due to their early experimentation with flavored nicotine vaping, such as increases in nicotine dependence.

For the first time since the 1990s, adolescent tobacco product use has increased, and the vast majority of adolescents are initiating nicotine use with flavored e-cigarettes (CDC, 2019). Although there is evidence from clinical and preclinical studies suggesting that flavorants increase nicotine reward in adult subjects, there is very little research on how e-cigarette flavorants affect nicotine reward and consumption during adolescence. Furthermore, we do not understand how initiation of nicotine with a palatable fruit-flavored product could affect long-term preferences for and consumption of nicotine as young vapers age. First-use of a flavored nicotine product could possibly increase the severity of nicotine addiction in a new generation of nicotine users.

This study was designed both to determine whether a fruit-flavorant can alter nicotine consumption in adolescent male and females, and to model the potential long-term consequences of initiating nicotine use with flavored products in adolescence.

## 2. Materials and methods

### 2.1 Animals

C57BL/6J mice were housed with a reverse 12 hr light/dark cycle (lights off at 10 A.M.) in a temperature-controlled room (24 ± 2 °C, relative humidity 55 ± 10%). Mice were weaned into single-housed cages with enrichment on post-natal day 21 (PND 21) with *ad libitum* access to food pellets (Labdiet 5001, PMI, Brentwood, MO) and to a source of liquid (see procedures below for more details). All acute behavioral testing occurred in the dark-phase, but drinking behavior was measured for 24-hours. Mice were evaluated in adolescence (PND 31-49), young adulthood (PND 50-70), and adulthood (PND 70+). Due to our interest in sex differences and the high “branching” nature of our longitudinal experimental, we estimated needing ∼75 mice to explore the possibility of sex differences at all phases of our experiment. This experiment was repeated in 4 separate groups of mice whose behavior was measured at different times throughout the year. Animals were bred in-house and were the offspring of 14 different breeding pairs. All procedures were approved by the institutional animal care and use committee and followed the guidelines for animal intramural research from the National Institute of Health.

### 2.2 Solutions

Both control and nicotine solutions contained 2% saccharin in filtered water, which was used to mask the bitter flavor of nicotine in nicotine-containing solutions (Perez et al., 2015; Salas et al., 2009; Zhang et al., 2012). For the majority of the experiment, nicotine bottles contained 0.1 mg/ml free-base nicotine (or 0.2804 mg/ml of nicotine hydrogen tartrate salt from Sigma Aldrich, St Louis, MO). During the “*Nicotine Fading + Flavor Reintroduction*” experiment, the nicotine concentration was sequentially reduced to 0.075, 0.05, and 0.025 mg/ml (free-base). Nicotine solutions were kept in dark bottles to protect against photodegradation and were prepared fresh every 4 days. Kool-Aid ® solutions were made according to package instructions (0.14 oz powder in 2 quarts of 2% saccharin solution). Kool-Aid ® powder (Kool-Aid ® Strawberry Drink Mix Unsweetened, Kraft Foods, Chicago IL) contains flavorants, but does not contain any additional sweeteners. In other words, the only sweetener in the Kool-Aid ® solutions was the saccharin that was added during preparation of the solution.

### 2.3 Strawberry vs. Grape Flavor Discrimination Test

Adult C57BL/6J mice of both sexes were tested for the ability to discriminate between Strawberry Kool-Aid ® solution and Grape Kool Aid Solution. Each mouse was transferred into a clean cage without food, water, or a nestlet, and was left to acclimate to this novel cage for 45 minutes. Following acclimation, 100 μl of water were added to the tip of a cotton swab and the swab was placed into the cage. The amount of time each mouse spent sniffing the cotton swab (oriented toward the swab and <2 cm away) was measured during a 2-minute odor presentation. A 1-minute break was given between each odor presentation. After three trials of water (control), mice were tested for time spent sniffing Strawberry Kool-Aid ®, and Grape Kool-Aid ® for three trials each. Light levels corresponded were between 0 and 2 Luxes in the testing area. Videos of each odor presentation were recorded and an individual blinded to treatment scored the time spent sniffing in each video.

### 2.4 Analyses of Kool-Aid ® Solution Volatiles

Headspace gas chromatography/mass spectrometry (GC/MS) analyses were conducted with a HT3 dynamic headspace analyzer (Teledyne Tekmar, Mason, OH, USA) outfitted with Supelco Trap K Vocarb 3000 thermal desorption trap (Sigma-Aldrich Co., St. Louis, MO, USA). The GC/MS (Thermo Scientific Trace Ultra) was equipped with a single quadrapole mass spectrometer (Thermo Scientific, Waltham, MA, USA) and a 30 m x 0.25 mm id Stabiliwax®-DA fused-silica capillary column (Restek, Bellefonte, PA, USA). Drinking solutions (2 mg/mL Kool-Aid ® in water without sweetener) were subjected to dynamic headspace GC/MS analysis by placing 50 μL in a sealed 20-mL headspace vial. The vial was maintained at 30 °C, swept with helium for 10 min (flow rate of 75 mL/min), and the volatiles collected on the thermal desorption trap. Trap contents were desorbed at 260 °C directly into the GC/MS using a split injection. The GC oven program had an initial temperature of 40 °C (held for 3.0 min) followed by a ramp of 7.0 °C/min to a final temperature of 230 °C (held for 6.0 min). The MS was used in scan mode from 33 to 400 m/z with a three minute solvent delay. Mass spectral peak identifications were assigned based on the library search of the NIST Standard Reference Database.

### 2.5 Measuring Longitudinal Flavor and Nicotine Preference and Consumption

The experiments described below were performed longitudinally. For a schematic of the experimental timeline see Figure 1. We utilized an adaption of the two bottle choice paradigm (2-BC) to evaluate the effect of flavorants on the voluntary consumption of nicotine. The volume of fluid consumed by animals was measured indirectly by weighing bottles (g) at each drinking time point. Assuming our solutions, which were made in filtered water, had a relative density of 1.0 g/mL, the loss of weight (g) from each bottle was considered to be equal to the volume consumed by the mouse in milliliters (mL). Mice were weighed daily during adolescence, and then every other day for the remainder of the experiment. The position of the bottles was alternated daily to account for possible side preferences. For each phase of the experiment, a control cage contained matching solutions, but no mouse. These control solutions were weighed to correct for solution loss that could have occurred due to evaporation and/or accidental movement of the housing rack. This control “drip volume” was subtracted from each animal’s “consumed volume” before preference and dose calculations were performed. Each phase in the experiment is described in detail below.

**Figure 1.**
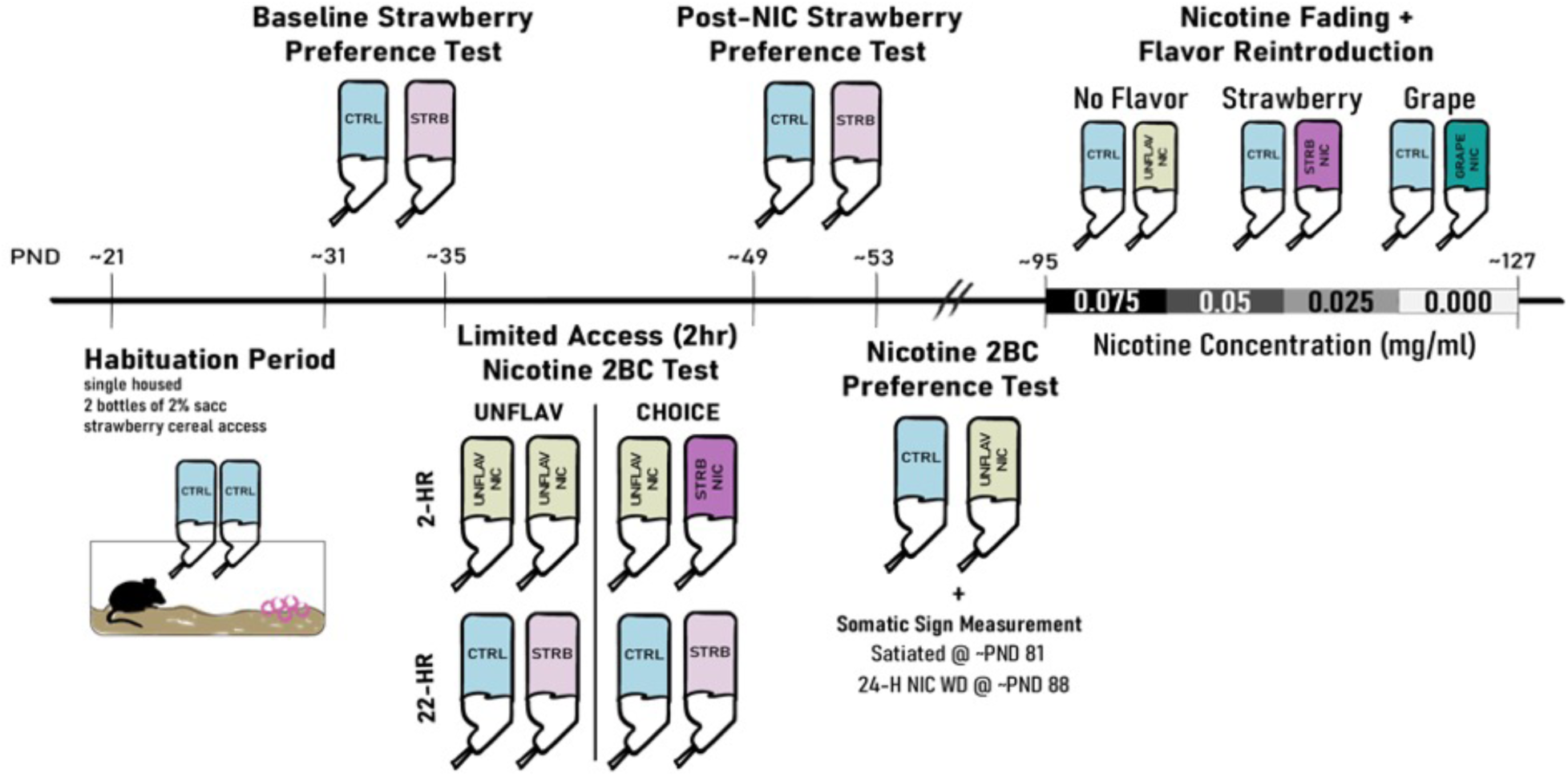
Timeline showing the sequence of events, drug treatments, and behavioral testing. Mice were divided into groups at two points in the experimental timeline. First, at the start of the “Limited Access (2hr) Nicotine 2 Bottle Choice (2BC) Test” mice were separated into either the “UNFLAV” or “CHOICE” groups. Data from the “Baseline Strawberry Preference Test” ensured that animals in each group had approximately equal baseline strawberry preference. All mice then followed the same treatment pattern, until “Nicotine Fading and Flavor Re-introduction”, when groups were further divided into one of three nicotine flavors: No Flavor, Strawberry Flavor, or Grape Flavor.

**Figure 2.**
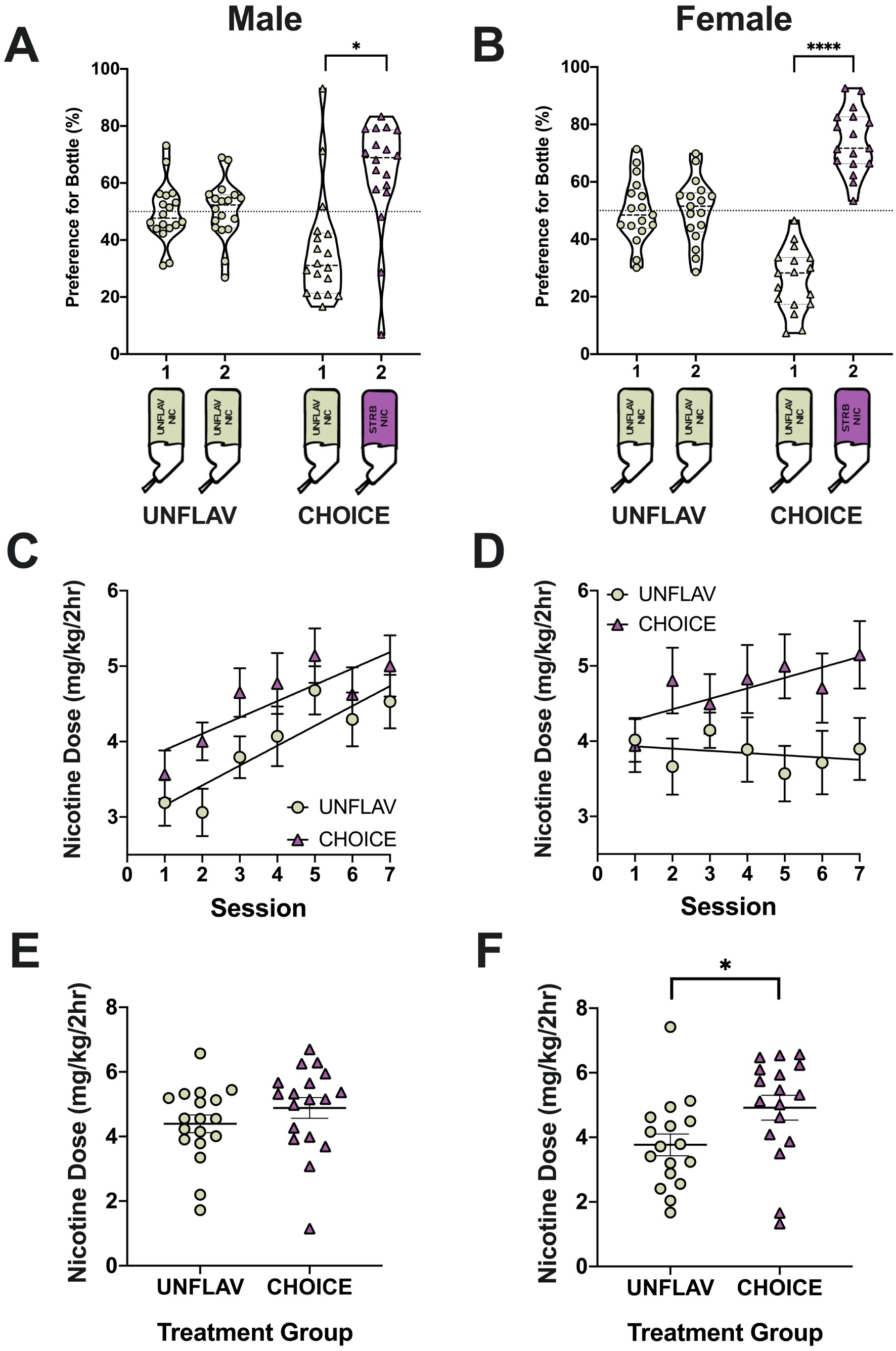
Adolescent mice of both sexes prefer a strawberry-flavored nicotine bottle in a two-bottle choice experiment, and females escalate nicotine consumption only if a strawberry-flavored nicotine solution is available. A-B) Violin plots display all data points of bottle preference in adolescence, with a thicker central representing the median and two thinner lines representing upper and lower quartiles. The violin plots show the preference for bottle 1 vs. bottle 2 in mice with access to two bottles of unflavored nicotine (UNFLAV) compared to mice with access to a bottle of unflavored nicotine (bottle 1) and a bottle of strawberry-flavored nicotine (bottle 2). C-D) Line graphs display the summary data (mean +/− SEM) of the total dose consumed by adolescent mice during a 2-hour nicotine access period of the course of 2-weeks of adolescence. A line of best-fit is overlayed for each data set. E-F) Display the average nicotine consumption during the last week of adolescence (Sessions 4-7). *P < 0.05 and ****P < 0.0001 for all comparisons. (Male, UNFLAV: n=18, CHOICE: n= 18; Female, UNFLAV: n=17, CHOICE: n=17).

**Figure 3.**
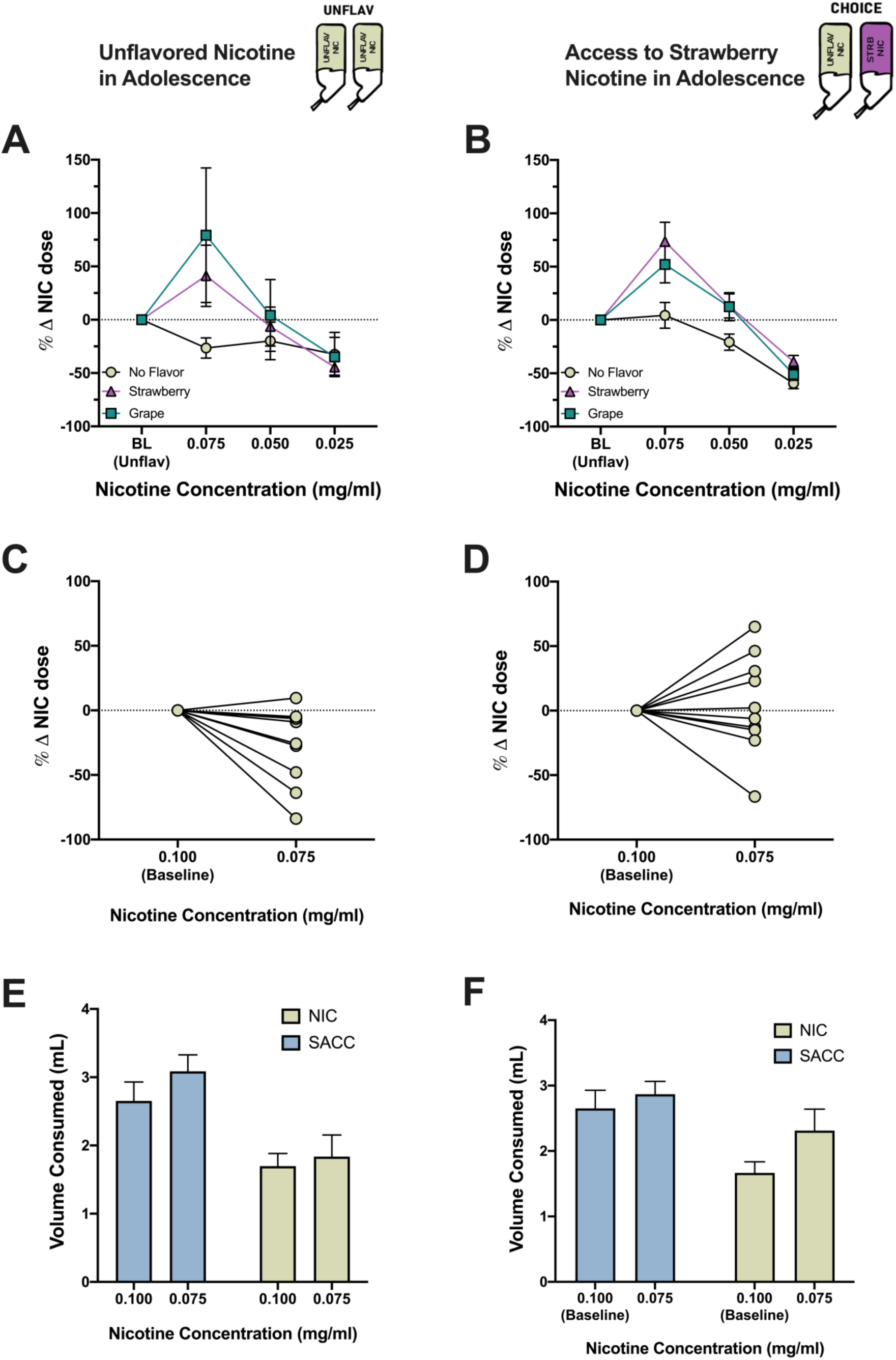
Adult mice increase nicotine consumption when palatable flavors are introduced into the bottle, and mice that initiated nicotine use with access to a palatable strawberry-flavored nicotine solution are resistant to a reduction in nicotine concentration. Data from mice that only had access to unflavored nicotine in adolescence (UNFLAV) (A,C,E) and mice that had access to strawberry flavored nicotine in adolescence (CHOICE) (B, D, F) are displayed on the left and right, respectively. In adulthood, the concentration of nicotine was reduced sequentially, and mice were either left with unflavored nicotine, had strawberry Kool-Aid® added to the nicotine bottle, or had Grape Kool-Aid® added to the nicotine bottle. A-B) Line graphs display the percent change of nicotine consumption (mg/kg) during the nicotine fading experiment relative to average baseline drinking (0.1 mg/ml nicotine solution) during maturation (mean +/− SEM). C-D) Before and after plots show the percent change of nicotine consumption from baseline (0.100 mg/ml nicotine solution) to 0.075 mg/ml nicotine solution, only in mice that had access to unflavored nicotine (i.e. ‘No Flavor’) during the fading experiment. E-F) Bar graphs show the volume consumed from the nicotine and saccharin bottles during the same period for mice that had access to unflavored nicotine (i.e. ‘No Flavor’) during the fading experiment.(UNFLAV in adolescence, ‘No Flavor’: n=12, ‘Strawberry’: n= 11, ‘Grape’: n=12; CHOICE in adolescence, ‘No Flavor’: n=10, ‘Strawberry’: n= 11, ‘Grape’: n=12).

#### 2.5.1 Habituation Period

On postnatal day (PND) 21 (±1 day), mice were single housed and given access to 2 bottles, each containing 2% saccharin in filtered water. Mice were also given 3 pieces of Strawberry Power O’s Cereal (Love Grown Goods, Denver, CO) daily, for 10 days (PND21-PND30). This was done to acclimate animals to strawberry flavoring in a naturalistic environment before experimentation with the flavor began.

#### 2.5.2 Baseline Strawberry Preference Test

After cereal feeding, mice began a two-bottle choice (2-BC) test (PND31), in which each mouse had access to a bottle containing 2% saccharin in filtered water and a strawberry-flavored bottle (Strawberry Kool-Aid ® in the same 2% saccharin solution). Bottles were weighed at 24-hour intervals for 4 days.

#### 2.5.3 Limited Access (2-hr) Nicotine Two-bottle Choice (2-BC) Test

For the remainder of adolescence (∼PND 35-50), mice were provided with 2 hours/day of nicotine access. Nicotine access began 2 hours after the start of the dark-phase (12 PM – 2 PM). The experimental group, referred to as the “CHOICE” group, had access to one bottle of unflavored nicotine (0.1 mg/ml nicotine (free-base) dissolved in 2% saccharin solution) and one bottle of strawberry-flavored nicotine (0.1 mg/ml nicotine (free-base) dissolved in Strawberry Kool-Aid ® + 2% saccharin solution). A control group, referred to as the “UNFLAV” group, had access to two bottles of unflavored nicotine. Mice were sorted into treatment groups so that they had approximately equal baseline strawberry preferences (Supplementary Figure 1). Experimenters weighed both nicotine bottles before and after the 2-hour session. Mice were excluded if they consumed less than 0.2 mL of nicotine during the 2-hr session on more than 2 days since this was similar to the average “drip” value measured in control-cage bottles. After the 2-hr drinking session, all animals were provided with 2 nicotine-free bottles for the remainder of the day (22-h). One of these bottles contained a 2% saccharin solution and the other contained a strawberry-flavored 2% saccharin solution.

#### 2.5.4 Post-Nicotine Strawberry Preference Test

24 hours after the last limited-access nicotine session (∼PND 51), mice were given one bottle of control solution (saccharin only) and one bottle of 0.1 mg/ml unflavored nicotine (free-base) + saccharin in the home cage. Access to these solutions continued for 96-h. This provided continued nicotine access, but allowed for a strawberry-flavor “wash-out” period before testing their post-nicotine strawberry preference. On ∼PND 55 mice began the ‘Post-Nicotine Strawberry Preference Test’, which was performed using an identical procedure to that detailed in the ‘baseline strawberry preference test’ section.

#### 2.5.5 Maturation Nicotine 2-BC

For the remainder of maturation (∼PND 59-95), mice had 24-h access to two bottles in the home cage. One bottle contained 0.1 mg/ml of free-base nicotine dissolved in a 2% saccharin solution and a second (control) bottle contained a 2% saccharin solution. Bottles were weighed at 24-h intervals. Drinking continued for ∼5 weeks.

#### 2.5.6 Spontaneous Withdrawal Testing

Approximately 3 weeks into the ‘Maturation Nicotine 2-BC’ phase, mice were observed while satiated with nicotine (i.e. at baseline) for the following physical signs: head shaking, scratching, grooming, chewing, jumping, paw licking, as described previously (Perez et al., 2015; Salas et al., 2009). Mice were then moved into a behavioral testing room with moderate lighting (Lux = 9-12) and allowed to acclimate for 1 hour. After acclimation, mice were placed in a new cage and their behavior was recorded on video for 20 min. One week later, the same procedure was repeated; however, to monitor signs of nicotine withdrawal, this time mice were recorded 24 hours after the start of nicotine deprivation. An individual blinded to animal ID, treatment, and state (e.g. satiated or withdrawn) measured the occurrence and the duration of the somatic signs. The total number of signs were used for comparison.

#### 2.5.7 Nicotine Fading and Flavor Reintroduction

At ∼PND 95, mice were sorted into 6 treatment groups based on “Adolescent Treatment” (e.g. CHOICE vs. UNFLAV) and an assigned “Fading Flavor” (e.g. ‘No Flavor’, ‘Strawberry’, or ‘Grape’). Mice were divided so that groups had approximately equal preference for strawberry at the end of adolescence, and approximately equal dose consumed-and preference for-nicotine during maturation (Supplementary Figure 4). During the “*Nicotine Fading and Flavor Reintroduction”* phase, mice had continued 24-hr access to one nicotine-containing bottle and one nicotine-free bottle, with two additional experimental manipulations. First, the concentration of nicotine in the nicotine-containing bottle was reduced, or “faded” from 1.0 mg/ml to 0.075 mg/ml, to 0.05 mg/ml, and finally to 0.025 mg/ml nicotine (free-base). Second, animals assigned to either a strawberry- or a grape-fading flavor had, in addition to a sequential reduction in nicotine concentration, their assigned fading flavorant introduced (or reintroduced, in the case of the CHOICE + ‘Strawberry’ fading flavor mice) into the nicotine bottle. All mice had a control bottle that contained a 2% saccharin solution. Exposure to each nicotine concentration lasted for 8 days.

**Figure 4.**
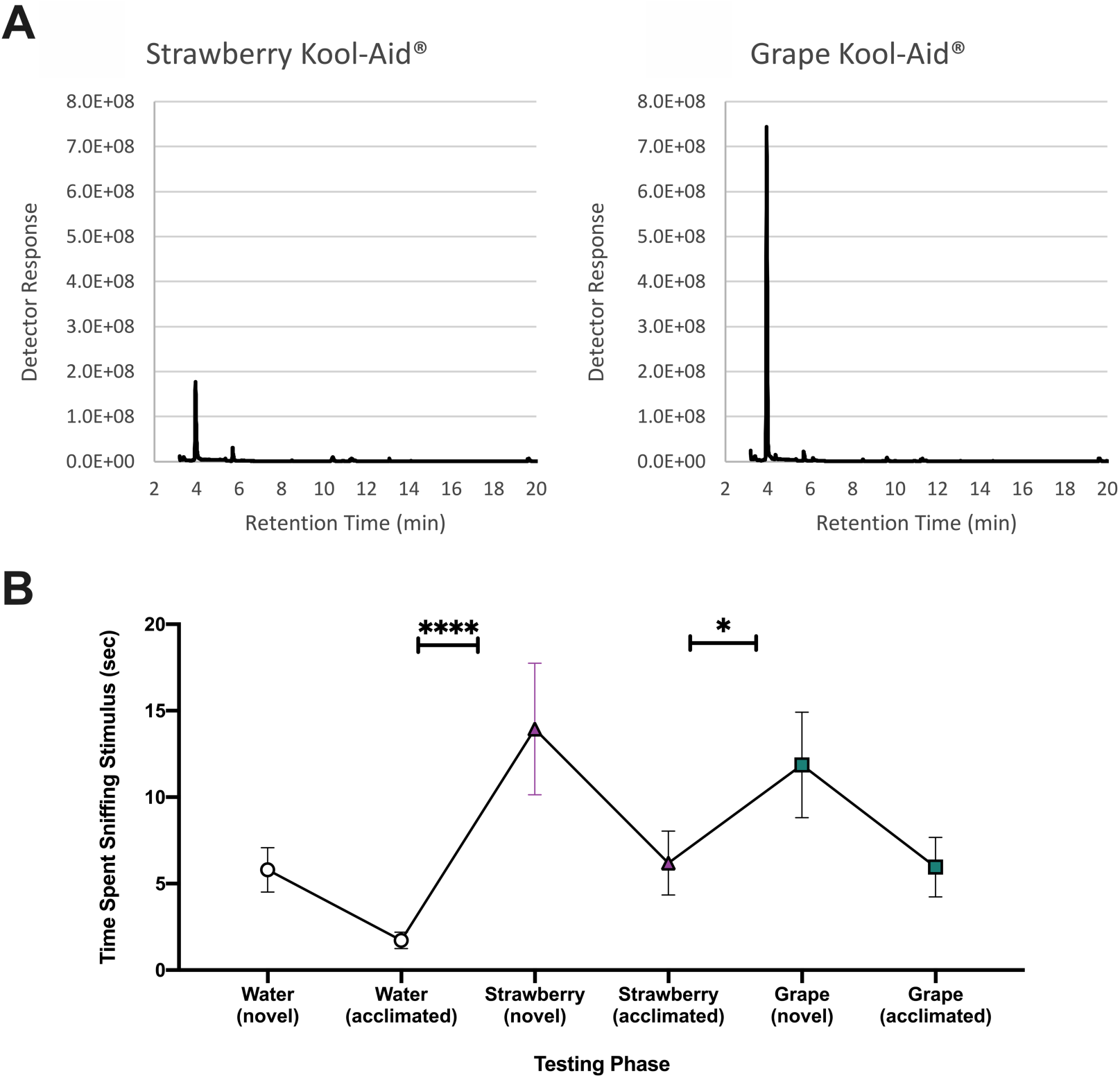
Mice discriminate between strawberry and grape Kool-Aid, despite the fact that both contain ethyl butyrate as the principal flavorant. (A) Chromatograms obtained from headspace gas chromatography/mass spectrometry (GC/MS) of aqueous solutions of strawberry (left) and grape (right) flavored Kool-Aid. The largest peak in each chromatogram (3.95 minutes) was identified as ethyl butyrate. (B) The line graph represents summary data (mean +/− SEM) of the time spent sniffing olfactory cues during an olfactory discrimination task. “Novel” data points represent the first exposure of a stimuli and “acclimated” data points are an average of the second and third trials with the same stimuli. (n= 19: 9 females, 10 males) *P < 0.05 and ****P < 0.0001 for all comparisons.

### 2.6 Data Analysis and Statistics

For drinking experiments, data were averaged over the course of two days (i.e. one session) so that each data point represents counterbalanced data in order to protect against misinterpretation due to innate side preferences of mice. For “Maturation 2-BC” data, where we observed mostly stable nicotine drinking, a within-animal outlier test (ROUT, Q=1%) was used to identify within-animal preference or consumption outliers. If a drinking session was identified as an outlier for an individual mouse that data point was replaced with the animal’s median data point during the corresponding phase of the experiment. All data sets were tested for normality using the D’Agostino Pearson Test prior to selecting appropriate methods of analysis (non-parametric vs. parametric). Non-parametric tests included simple linear regression, Wilcoxon sign rank test, Friedman’s test, and Generalized Estimating Equations (GEE) analyses. Parametric tests employed in this study include t-tests, paired t-tests, unpaired t-tests, and fixed effects ANOVA and RMANOVA. Data are represented as +/− SEM for all data. GraphPad Prism (San Diego, CA) was used for the majority of data analyses, with the exception of the Generalized Estimating Equation analysis which was run using SPSS.

## 3. Results

### 3.1 Adolescent mice prefer a strawberry-flavored nicotine solution over an unflavored nicotine solution

Strawberry-flavored (vs. unflavored) nicotine preference was measured in adolescent mice during the “*Limited Access (2-hr) Nicotine Two-bottle Choice (2-BC) Test”* (see Figure 1 for experimental schematic). Mice were divided in two experimental groups: one group had access to two bottles of unflavored nicotine (i.e. “UNFLAV” group) and a second group had access to one bottle of unflavored nicotine and one bottle of strawberry-flavored nicotine (i.e. “CHOICE” group) during a 2-hour nicotine access period. An initial two-way ANOVA of CHOICE mice bottle preference with factors of sex and bottle detected a significant effect of sex (F 1,33)_sex_ = 48.23, ****p <0.0001). As a result, male and female data is displayed and analyzed separately. Female adolescent data show an effect of both bottle and interaction between bottle and treatment group (two-way ANOVA, F_bottle_(1,32) = 40.12, ****p<0.0001, F_interaction_(1,32) = 39.61, ****p<0.0001), with Sidak’s multiple comparison’s test detecting female CHOICE mice have a significant preference for the strawberry-flavored nicotine bottle (****p<0.0001) (Figure 2B). Male CHOICE mice also show a significantly greater preference for the strawberry-flavored nicotine bottle (Wilcoxon matched-pairs signed rank test, n=18, *p=0.01) (Figure 2A). No statistically significant preference for either bottle was seen in mice in the UNFLAV group (Female: Sidak’s multiple comparison’s test, p = 0.9995, Male: paired t-test, t(17) = 0.1889, p=0.8524). (Figure 2A-B).

### 3.2 Female mice only escalate nicotine consumption during adolescence if a strawberry-flavored nicotine solution is provided

Males in both the CHOICE group and the UNFLAV group escalated nicotine consumption over the course of adolescence (Figure 2C). Escalation is indicated by a significant increase in nicotine consumption between session 1 and session 7 (UNFLAV male mice, paired t-test, t(17)=3.449, **p = 0.0031; CHOICE male mice, paired t-test, t(17)=3.081, **p=0.0068). However, while female mice with access to strawberry-flavored nicotine escalate nicotine consumption (paired t-test, t(16)=2.544, *p=0.0216), while the females in the UNFLAV group maintained their dose without escalation (paired t-test, t(16) =0.2478, p = 0.8074) (Figure 2D).

### 3.3 Adolescent female mice drink a higher dose of nicotine when given access to a strawberry-flavored nicotine solution

The total amount of nicotine consumed during adolescence (measured in mg/kg body weight) was measured following each 2-hour drinking period of the “*Limited Access (2-hr) Nicotine Two-bottle Choice (2-BC) Test”*. On average, during the last week of adolescence (sessions 4-7) male mice in the CHOICE group (i.e. mice with access to strawberry-flavored nicotine) drank 11.2% more nicotine (4.885 ± 0.318 mg/kg) than their counterparts in the UNFLAV group (4.393 ± 0.275 mg/kg) (Figure 2E). Whereas, female mice in the CHOICE group drank 30.5% more nicotine (4.918 ± 0.386 mg/kg) on average during the last week of adolescence than their counterparts in the UNFLAV group (3.768 ± 0.335 mg/kg), which rose to the level of significance (unpaired t-test, t(32) = 2.251, *p=0.031) (Figure 2F). Although mice in the CHOICE group tend to drink a significantly greater volume of fluid during the 2-hour nicotine session resulting in higher consumed nicotine doses, the volume of fluid consumed by animals on average during 24-hours does not differ (Supplementary Figure 2).

### 3.4 Mice that initiate nicotine consumption in adolescence with a fruit-flavorant resist reductions in nicotine consumption in adulthood

The same mice that were used in the “*Limited Access (2-hr) Nicotine Two-bottle Choice (2-BC) Test”* were allowed to mature with access to one bottle containing 2% saccharin and one bottle containing 0.1 mg/ml nicotine in 2% saccharin during the *“Maturation Nicotine 2-BC”* phase (see Figure 1 for experimental schematic). Adolescent treatment (i.e. UNFLAV or CHOICE) had no effect on preference for- or dose consumed of-nicotine during this maturation phase, when mice had access to 0.10 mg/ml unflavored nicotine (free base) 24h/day (Supplementary Figure 3). However, differences based on adolescent treatment emerged when the nicotine concentration was decreased over time in the nicotine bottle. A preliminary Generalized Estimating Equations (GEE) analyses that included sex as a factor did not detect any sex effects, so data were pooled and followed up with GEE analyses considering the following factors: adolescent treatment, nicotine concentration, fading flavor, and all associated interactions. Parameter estimates of the GEE analyses indicate a significant effect of fading flavor (**p=0.001), nicotine concentration (****p<0.0001) and a significant interaction between fading flavor *nicotine concentration (****p<0.0001). However, if a palatable flavor was not added to the bottle (i.e. unflavored nicotine during fading), consumption dropped below baseline (Figure 3A). Mice that had not been exposed to flavored nicotine during adolescence (UNFLAV group) and that received progressively diluted unflavored nicotine solutions during the “*Nicotine Fading and Flavor Reintroduction*” phase, reduced their nicotine consumption when the nicotine concentration dropped to 0.075 mg/ml. (Figure 3A and C). Conversely, mice exposed to flavored nicotine during adolescence (CHOICE group) compensated for the drop in the concentration of unflavored nicotine from 0.1 to 0.0075 mg/ml by increasing the volumes drank, thereby maintaining the nicotine dose to pre-fading levels for an additional 8 days compared to the UNFLAV group (Figure 3B, D, and F). Following the GEE analysis and considering the fact that all ‘No Flavor’ fading data were normally distributed, the response of mice in this treatment group to the initial reduction in nicotine concentration from 0.10 mg/ml nicotine to 0.075 mg/ml was further analyzed using parametric testing. Both an effect of adolescent treatment (two-way RMANOVA, F_adolescenttreatment_(1,20) = 4.088, p=0.0568) and an interaction between adolescent treatment and nicotine concentration (F_interaction_(1,20) = 4.088, p=0.0568) fell just short of significance. In order to determine if this difference in dose-maintenance was related to differences in nicotine dependence, we measured the number of somatic behaviors (e.g. grooming, chewing, licking, etc.) displayed by animals in videos taken during both during nicotine satiety and 24-hours into nicotine withdrawal. Although on average mice did not display significant signs of spontaneous nicotine withdrawal (Supplementary Figure 3D), there was a trend towards an increase in the proportion ‘CHOICE’ mice that displayed symptoms of nicotine withdrawal (defined by > 15% increase in the number of somatic signs performed) (Supplementary Figure 3E).

### 3.5 Mice increase nicotine consumption if a palatable fruit-flavorant is added to nicotine solutions in adulthood

We next considered the effect of adding palatable flavors, both familiar (strawberry) and unfamiliar (grape) would have on nicotine consumption during adulthood. Interestingly, both the UNFLAV and CHOICE groups increased their drinking behavior when presented with diluted – but flavored-nicotine. This increase occurred despite the reduction in nicotine concentration from 0.1 mg/ml to 0.075 mg/ml free-base nicotine (Figure 3A & B). This effect was not specific to the familiar flavorant (i.e. strawberry), as it was also observed when a novel fruit flavorant (i.e. grape) was added to the nicotine solution. To understand the potential mechanisms for such response, we first assessed the chemical composition of strawberry- and grape-flavored Kool Aid® flavorants by GC/MS (Figure 4A & B). Although the two fruit flavorants share a major chemical component in ethyl butyrate (Figure 4A), mice can discriminate between Strawberry and Grape Kool-Aid® solutions in an olfactory discrimination task, as indicated by an increase in time spent sniffing Strawberry Kool-Aid® at first presentation (****p<0.0001, Friedman test followed by Dunn’s multiple comparison test) and an increase in sniffing grape Kool-Aid® when presented following acclimation to strawberry Kool-Aid (*p=0.0431, Friedman test followed by Dunn’s multiple comparison test) (Figure 4C). The fact that mice identify these solutions as unique, despite containing the same major chemical constituent, suggests that ethyl butyrate is not indiscriminately increasing nicotine intake, but more likely is contributing to a unique sensory profile of the flavorants, which ultimately promotes consumption via secondary factors.

## 4. Discussion

The potential of flavored e-cigarettes to increase adolescent nicotine reward and consumption has been a major public health concern. In addition, individuals who report a positive first-experience with smoking are more likely to go on to become regular smokers (Chen et al., 2003; DiFranza et al., 2007; Mantey et al., 2017; Rodriguez and Audrain-McGovern, 2004; Sartor et al., 2010; Urbán, 2010). Therefore, it is important to consider how initiating nicotine use with palatable flavored e-cigarettes could impact long term nicotine consumption and reward. In this study, we investigated the influence that fruit flavorants could have on nicotine consumption in adolescence and we examined how this “palatable/positive” first nicotine exposure might impact later nicotine consumption and preference with and without flavorants. To examine the abuse liability of flavored nicotine long term, from adolescence to adulthood, we chose the 2BC model of voluntary oral consumption and preference. Compared to the intravenous self-administration paradigm, this approach allowed us to overcome the challenges associated with the small size of young mice and the need for prolonged catheter patency. Despite resulting in slower pharmacokinetics compared to intravenous nicotine self-administration, the 2BC model produces an expected inverted U-shape dose-response curve for nicotine and leads to symptoms of withdrawal when nicotine is withheld (Bagdas et al., 2019; Pogun and C Collins, 2012).

Both male and female adolescent mice prefer a nicotine solution when it was flavored with a palatable fruit flavorant (i.e. strawberry); however, this preference was more prominent in female mice. Access to flavored nicotine also had a unique effect on adolescent female nicotine consumption. Whereas male mice escalated nicotine consumption over the course of adolescence regardless of their access to flavored nicotine, female mice *only* escalated nicotine consumption if they had access to the palatable strawberry-flavored nicotine solution. Evidence from human literature supports the idea of sex differences in relation to perceived palatably of flavored tobacco products. Women are more likely than men to use characterizing flavored products and to value flavor availability (Kistler et al., 2017; Patten and De Biasi, 2020; Perkins, 1999; Perkins et al., 2001; Soneji et al., 2019; Xiao et al., 2019). Furthermore, there is evidence that women are more driven by the sensory aspect of smoking, as opposed to the pharmacological effects of nicotine (Perkins, 1999; Perkins et al., 2001a; US Surgeon General, 2001). It is therefore possible that adolescent female mice drank significantly more nicotine when they had access to flavored-nicotine because they were driven by the ‘flavor’ and less so, by nicotine. Regardless of their motivation, the fact that female mice drank ∼30.5% more nicotine when they had access to flavored nicotine (compared to males who drank ∼11.2% more) has major implications for young women’s health and development. Adolescence is a period of sensitive neurodevelopment and nicotine exposure during adolescence is linked to a multitude of health complications, including cognitive deficits and increased probability of becoming a dependent smoker in adulthood (England et al., 2017, 2015; Goriounova and Mansvelder, 2012; US HHS, 2016; Omelchenko et al., 2016; Walker and Loprinzi, 2014; Yuan et al., 2015).

A recent meta-analysis showed a strong and consistent association between initial e-cigarette use and subsequent cigarette smoking initiation (Soneji et al., 2017). In addition, flavored e-cigarette use (rather than e-cigarette use generally) has been associated with higher rates of vaping and increased risk of subsequent cigarette initiation (Barrington-Trimis et al., 2018; Chen et al., 2016; Dai and Hao, 2016; Leventhal et al., 2019b). Therefore, we were interested in the possibility that mice with access to flavored nicotine in adolescence (i.e. CHOICE mice) would have higher preferences for nicotine and consume higher doses of nicotine during maturation. Our data did not indicate an increase in nicotine preference or consumption during young adulthood, once flavor was removed from the bottle. Interestingly, not all research suggests that flavors are associated with a progression to combustible use (Audrain-McGovern et al., 2019). Furthermore, CDC data published in 2019 only shows an increase in e-cigarette, not combustible smoking, among adolescent populations (CDC, 2019). Therefore, the role of flavored e-cigarettes on the ability to promote progression to combustible cigarettes is still unclear.

Although adolescent access to fruit-flavored nicotine did not alter nicotine consumption at a stable nicotine concentration during adulthood, when the concentration of the nicotine solution was reduced by 25%, only mice from the CHOICE group maintained baseline nicotine consumption. It is important to note that mice from both the CHOICE and UNFLAV adolescent treatment groups had similar preferences for nicotine and consumed similar doses of nicotine during the maturation phase. Neither group showed evidence of nicotine withdrawal, when physical signs during withdrawal were compared to those measured during the nicotine-sated state. This is possibly due to the fact that spontaneous withdrawal is, in general, more difficult to quantify than pharmacologically precipitated withdrawal. There are other mechanisms through which the adolescent strawberry nicotine exposure could have altered the value of nicotine to animals in adulthood. For example, adolescent strawberry nicotine exposure could have strengthened the CHOICE mice’s secondary dependence motives (SDMs) for nicotine consumption. SDMs include ancillary factors regulating nicotine dependence like affiliative attachment, cue exposure, and taste, and are strongly correlated to pleasure and relief derived from nicotine use (Smith et al., 2010; Tarantola et al., 2017). In contrast, primary dependence motives (PDMs) are the core features of nicotine dependence including factors like automaticity, craving and tolerance that are more related to physical nicotine dependence (Adkison et al., 2016; Smith et al., 2010). Score on an SDM scale is less correlated with nicotine dependence than PDMs and is not correlated with daily cigarette count, but is still a significant predictor of cigarette craving (Tarantola et al., 2017). Menthol cigarette smokers show increased affiliative attachment to cigarettes compared to non-flavored cigarette smokers, illustrating the possibility of nicotine flavorants to affect SDMs (Cwalina et al., 2020). Therefore, it is possible that our CHOICE group maintains baseline nicotine dose during tapering due to increased SDMs causing increased craving without affecting physical withdrawal or baseline nicotine consumption, although further research would be necessary to identify the specific behavioral features underlying this finding.

For all mice, regardless of prior exposure to flavored nicotine, the addition of either strawberry (the familiar flavorant) or grape (a novel fruit flavorant) led to a 25-100X increase in nicotine consumption, despite a 25% decrease in nicotine concentration. The increase in preference for, and consumption of fruit-flavored nicotine in adolescence and adulthood observed in this study could be motivated by either an increase in nicotine reward or a ‘masking’ of the harshness of nicotine. Tobacco company documents openly state that inclusion of sweet and fruity flavor additives (e.g. “vanilla beans, peach, apricot, licorice, cocoa, and many others”) are meant to cover up “objectionable off flavors” (Fries and Brother and Triest; Kostygina et al., 2014; Industry document, 1966) and the addition of characterizing flavors can suppress ‘unappealing’ sensations of nicotine in e-cigarettes and of other bittering agents (Isogai and Wise, 2016; Leventhal et al., 2019a). Therefore, it is possible that in our study, flavorants overpowered the bitterness in their nicotine solutions and led to increased consumption of the more palatable solution. On the other hand, it has been shown that fruity flavorants are able to increase ratings of sweetness (Frank and Byram, 1988; Labbe et al., 2007; Rao et al., 2018; Smith et al., 2020). Mass spectrometry of Kool-Aid ® solutions revealed that the major component of both the strawberry and grape Kool-Aid ® solutions was ethyl butyrate. Ethyl butyrate (also known as ethyl butanoate) was recently determined to be the second most common flavorant in a survey of 277 e-cigarette refill liquids (Omaiye et al., 2019) and has been shown to increase ratings of the ‘sweetness’ of solutions (Rao et al., 2018). ‘Sweetness’ is naturally rewarding and therefore could work to enhance nicotine reward. Considering that both strawberry and grape flavorants were able to interfere with nicotine devaluation in the nicotine fading paradigm, our results might be explained by the presence of ethyl butyrate in both nicotine solutions. More research would be needed to determine which of these factors was the driving force behind the increase in nicotine consumption we observed.

One of the few pre-clinical studies to-date which investigated the role of a fruit flavorant on nicotine reward saw increased conditioned place preference in adult male mice that were injected with the ‘green apple’ flavor (e.g. farsenol) in combination with nicotine (Avelar et al., 2019). This report was remarkable in that it suggests that e-cigarette flavor additives have pharmacological effects of their own. However, since flavorants were injected, the study could not account for the potential sensory contributions of the fruit flavorant. Interestingly, Avelar et al. found that farsenol only enhanced nicotine reward in male mice, whereas, in this study we saw that strawberry Kool-Aid ® enhanced nicotine consumption more drastically in females. This could indicate that each flavorant (e.g. farsenol vs. ethyl butyrate) will have unique sex-dependent effects on nicotine reward and/or consumption. Alternatively, it is possible that the sensory components of flavorants are necessary to fully appreciate the effect of flavorants on nicotine reward, particularly among females. In fact, a recent publication also found that fruit, but not tobacco flavored e-liquids enhanced nicotine consumption in adult male mice (Wong et al., 2020). Despite this increase in oral nicotine consumption, they detected no enhancement of nicotine conditioned place preference when e-cig flavorants were injected intraperitoneally, suggesting that in that case, orosensory factors and not pharmacological factors enhanced voluntary nicotine consumption. Studying e-cigarette flavorants in oral administration paradigms, as was done by Wong and colleagues and here, is better suited to contribute to our understanding of the sensory impact of flavors on nicotine reward and consumption. Ultimately, an understanding of both sensory and pharmacological contributions is necessary to decipher how the availability of flavorants might impact nicotine use.

This research highlights the importance of studying flavored nicotine using techniques that allow for the sensory components of flavorants to be perceived by research subjects, as well as the importance of studying both male and female subjects. One weakness to our experimental design is that we are unable to distinguish between changes in nicotine consumption due to increased reward or reduced aversion of nicotine. Future studies which combine both pharmacological and sensory components of popular e-cigarette flavorants, such as via vapor inhalation, might help address these questions. Nevertheless, our results indicate that access to flavored nicotine leads to increased consumption during adolescence, particularly among females. This could lead to consequences ranging from long-term cognitive effects to an increased likelihood of being a dependent smoker in adulthood. We saw no evidence that first exposure to a flavored nicotine product impacted nicotine preference or consumption as an adult until nicotine concentration was reduced, at which point mice that initiated with strawberry-flavored nicotine resisted reductions in nicotine dose. As mentioned earlier, the mechanism for this behavior remains unclear since no differences in nicotine preference, consumption, or dependence were measured. However, it could have interesting implications on future difficulties during nicotine cessation attempts by individuals who initiate with a flavored product in adolescence.

## 5. Conclusion

In summary, our results indicate that fruit-flavorants have a significant impact on nicotine preference and consumption, having the ability to increase nicotine consumption, regardless of life-stage and could potentially interfere with one’s ability to quit later in life after exposure to flavored nicotine during adolescence.

## Acknowledgements

The authors would like to acknowledge Kimberly Halberstadter for her help with the bottle choice experiments and Annie Luo for scoring the odor discrimination videos.

## CRediT Author Statement

**Theresa Patten:** Conceptualization, Data curation, Formal analysis, Investigation, Methodology, Supervision, Writing - original draft, Writing - review & editing **Allison Dreier:** Conceptualization, Data curation, Investigation, Methodology **Rae J. Herman**: Data curation, Writing - review & editing **Bruce A. Kimball**: Data curation, Writing - review & editing **Mariella De Biasi**: Conceptualization, Funding acquisition, Project administration, Resources, Supervision, Writing - review & editing.

