## supplementary figures for "Exposure to fruit-flavoring during adolescence increases nicotine consumption and promotes dose escalation"

**Supplementary Figure 1.**

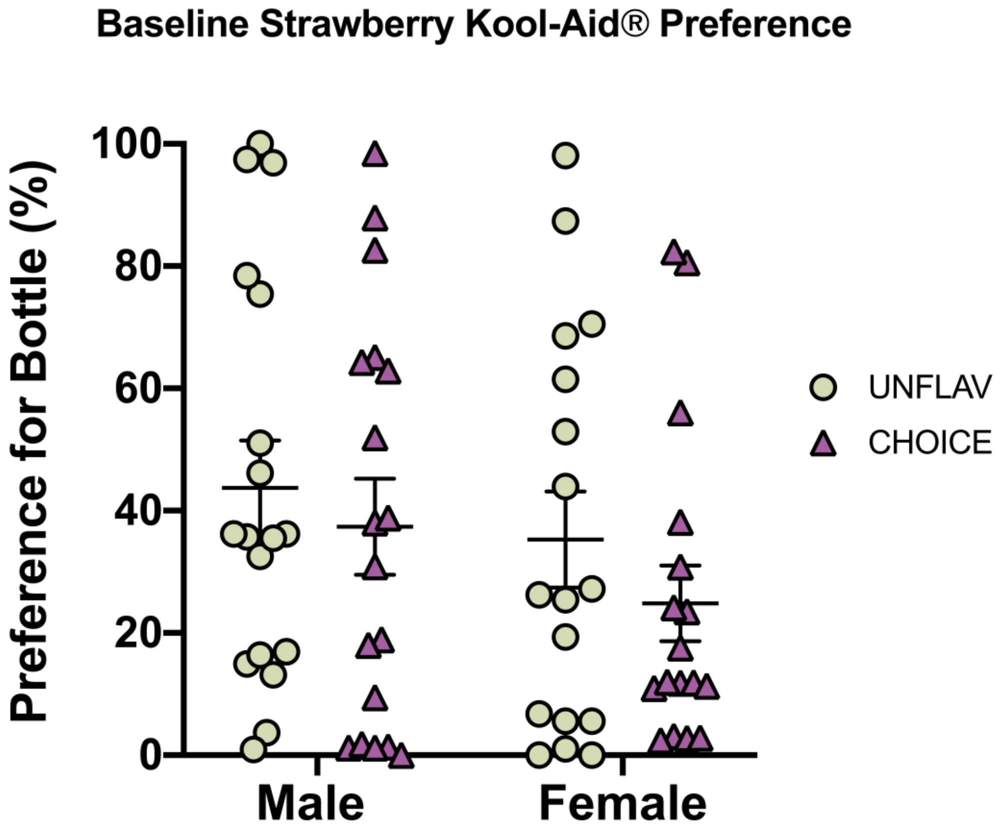


**Animals are grouped such that strawberry preference is balanced across treatment groups (i.e. UNFLAV, CHOICE).** Following cereal pre-treatment, mice are given a "*Baseline Strawberry Preference Test"* in which the volume consumed by mice from a strawberry-flavored bottle and from an unflavored bottle are measured at 24-hour intervals. Mice were assigned into treatments groups such that there is no difference in preference between mice sorted into the UNFLAV group (baseline strawberry preference = 39.64 ± 5.47) and those sorted into the CHOICE group (baseline strawberry preference = 31.29 ± 5.08). Lines and error bars represent summary data (mean +/- SEM). Two-way ANOVA did not detect a significant effect of sex, treatment, or an interaction ($F_{sex}(1,66) = 1.980$, $p=0.164$, $F_{treatment}(1,66) = 1.271$, $p=0.263$, $F_{interaction}(1,66) = 0.075$, $p=0.784$). (Male, UNFLAV: n=18, CHOICE: n= 18; Female, UNFLAV: n=17, CHOICE: n=17).

**Supplementary Figure 2.**

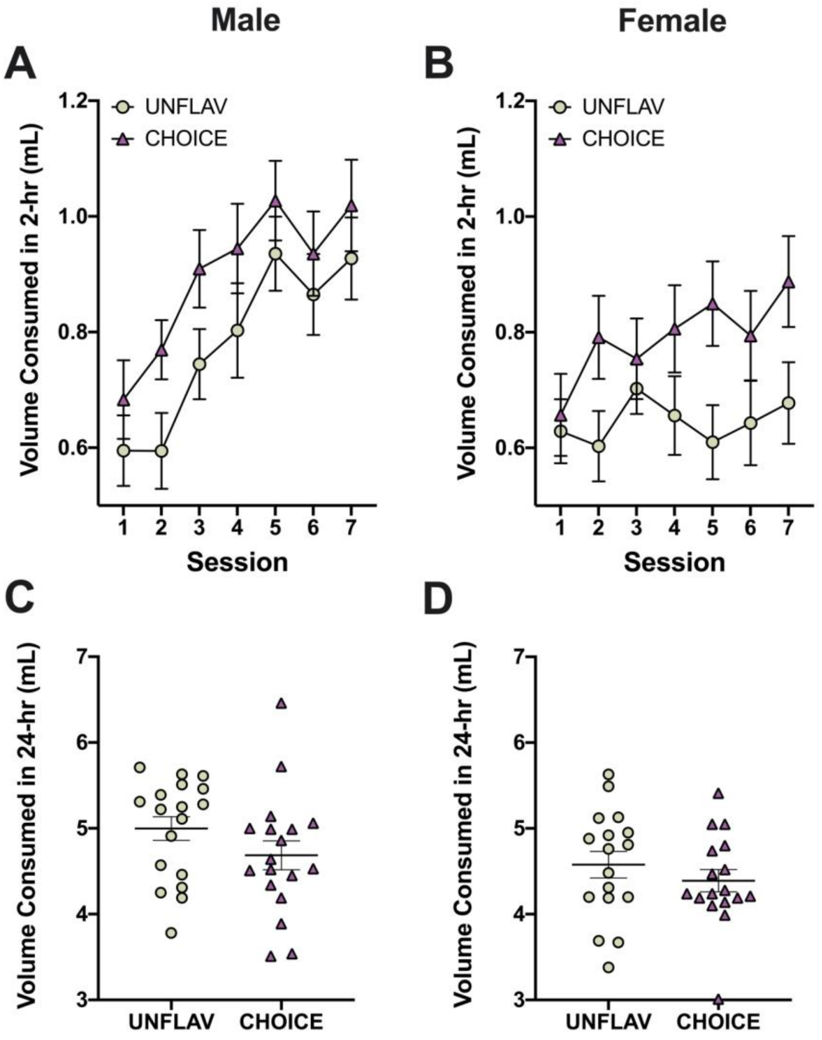


**Mice with access to strawberry flavored nicotine tend to drink a larger volume of nicotine during the 2-hour nicotine access session, but do not drink more fluid overall.** A-B) The summary data of total volume (mL) consumed during the 2-hour nicotine access period in adolescence is shown for males and females separately (mean +/- SEM). A 2-way RMANOVA (treatment x session) detected a trend but no significant effect of treatment group on the volume consumed during the 2-hour nicotine session among males ($F_{group}(1,34) = 2.868$, $p=0.100$) and B) among females ($F_{group}(1,32) = 3.676$, $p=0.064$). There was a significant effect of session for males ($F_{session}(6,204)=13.22$, ****$p<0.0001$), but not females ($F_{session}(6,192)=1.664$, $p=0.132$), and no significant interactions (Male: $F_{interaction}(6,204)=0.3227$, $p= 0.925$; Female: $F_{interaction}(6,192)=1.460$, $p=0.194$). C-D) Data shows the average volume consumed by mice over 24-hr. Lines and error bars represent summary data (mean +/- SEM). The volume consumed over a 24-hr period is not significantly different between treatment groups in either C) male (unpaired t-test, $t(34) = 1.428$, $p=0.162$) or D) female mice (unpaired t-test, $t(32) = 0.924$, $p=0.3623$). (Male, UNFLAV: n=18, CHOICE: n= 18; Female, UNFLAV: n=17, CHOICE: n=17).

**Supplementary Figure 3.**

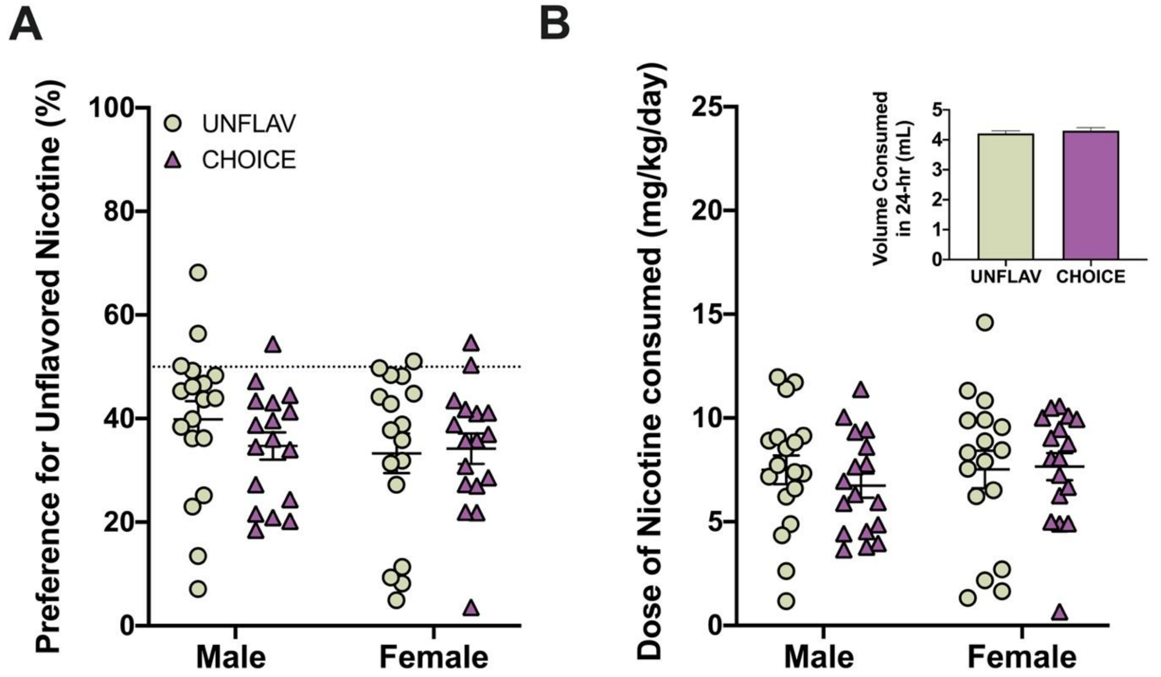


**Access to strawberry-flavored nicotine in adolescence does not increase preference for or consumption of unflavored nicotine during maturation.** Following the "Limited Access (2-hr) Nicotine 2BC Test" in adolescence, all mice were given one bottle containing 0.1 mg/ml free base nicotine and 2% saccharin and one bottle that contained 2% saccharin during the "Maturation Nicotine 2-BC" phase. Neither adolescent treatment (e.g. UNFLAV or CHOICE), sex, nor an interaction affected A) the preference for an unflavored nicotine solution during maturation (2-way ANOVA, $F_{sex}(1,65) = 1.182$, p=0.281, $F_{treatment}(1,65) = 0.429$, p=0.515, $F_{interaction}(1,65) = 0.857$, p=0.358). B) Adolescent treatment, sex, nor an interaction affect also does not affect the dose consumed (mg/kg) of an unflavored nicotine solution during maturation (2-way ANOVA, $F_{sex}(1,65) = 0.425$, p=0.517, $F_{treatment}(1,65) = 0.189$, p=0.665, $F_{interaction}(1,65) = 0.393$, p=0.533). Lines and error bars represent summary data (mean +/- SEM). The volume consumed by mice of both sexes during 24-hour was averaged across all maturation drinking sessions and no difference was detected (unpaired t-test, t(67)=0.6443, p=0.52). (Male, UNFLAV: n=18, CHOICE: n= 18; Female, UNFLAV: n=17, CHOICE: n=17).

**Supplementary Figure 4**

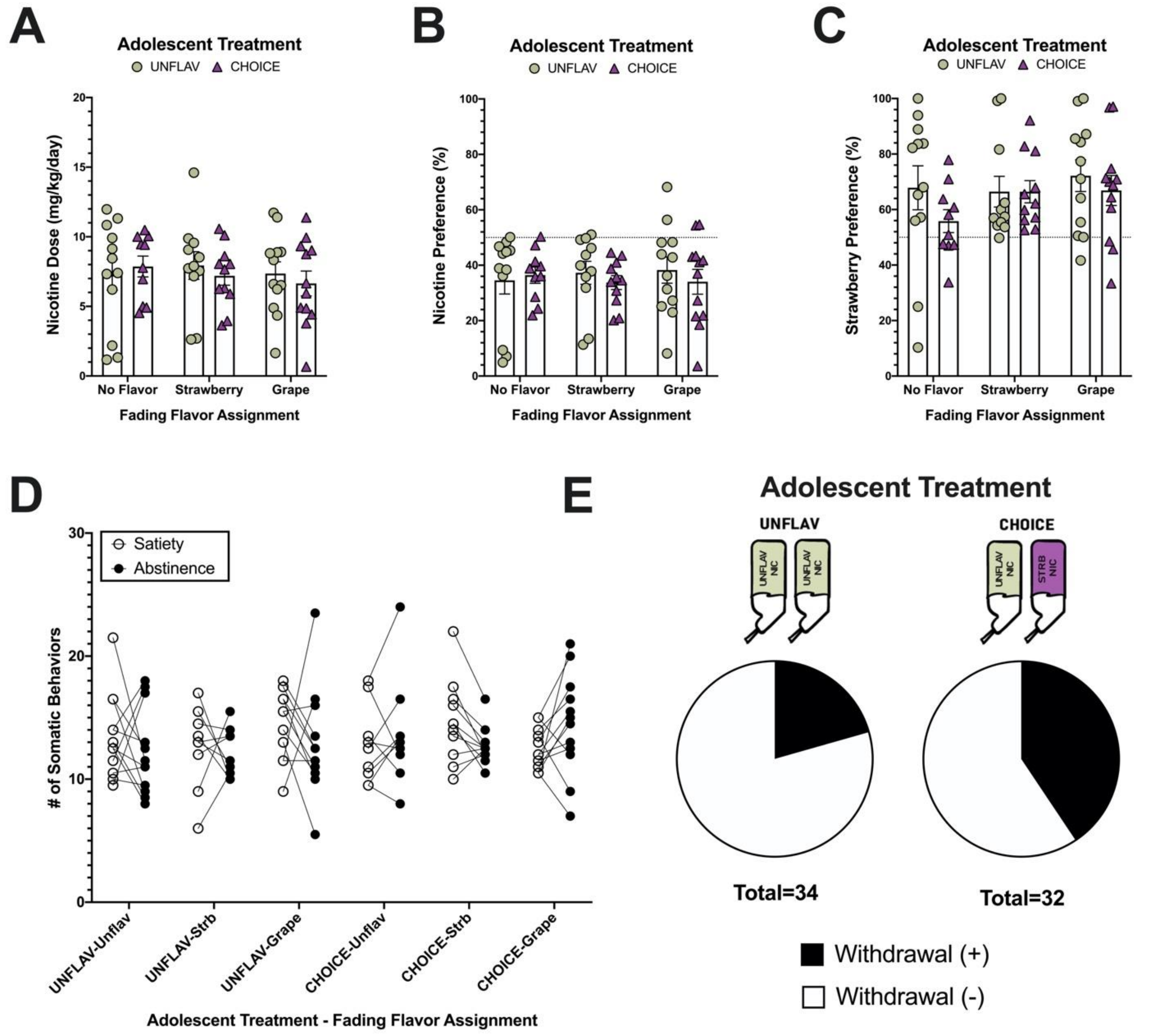


**Mice from each fading group have similar preferences for nicotine and strawberry solutions, consume similar doses of nicotine, and do not exhibit significant signs of nicotine dependence at the start of the "Fading Nicotine Concentration with Flavors 2-BC" experiment.** Bar graphs represent summary data (mean +/- SEM). A) The average dose consumed by mice during maturation was 7.34 $\pm$ 0.36 mg/kg with no effect of adolescent treatment (two-way ANOVA, $F_{adolescent\ treatment}(1,62) = 0.1522$, p=0.70) and with animals assigned into groups such that no statistical differences were observed in any group ($F_{fading\ group}(2,62)$ = 0.2696, p=0.76). B) The average preference for a 0.1 mg/ml nicotine bottle compared to control was 35.69 $\pm$ 1.65 with no effect of adolescent treatment (two-way ANOVA, $F_{adolescent\ treatment}(1,62)$ = 0.3296, p= 0.57) and with animals assigned into groups such that no statistical differences were observed in any group ($F_{fading\ group}(2, 62)$ = 0.01649, p= 0.9837). C) The average preference for a strawberry-flavored solution compared to control during the "Post-NIC Strawberry Preference Test" was 66.21 $\pm$ 2.33 with no effect of adolescent treatment (two-way ANOVA, $F_{adolescent\ treatment}(1,62)$ = 1.521, p = 0.2221) and with animals assigned into groups such that no statistical differences were observed in any group ($F_{fading\ group}(2, 62)$ = 0.9122, p= 0.4069). D) The number of somatic signs displayed by animals during two trials (satiated vs. 24-hr nicotine abstinent) were compared. A two-way RMANOVA did not detect a significant effect of state (i.e. satiety vs. abstinent) ($F_{state}(1, 60)$ = 0.7993, p = 0.3749) signifying that, on average, the 4 weeklong 24-h nicotine choice was not sufficient to produce nicotine dependence in these animals. E) There is however, a trend towards an increase in the proportion 'CHOICE' mice that displayed symptoms of nicotine withdrawal (defined by $\geq$ 15% increase in the number of somatic signs performed) compared to 'UNFLAV' mice (Fisher's exact test, p=0.109)

**Supplementary Figure 5.**

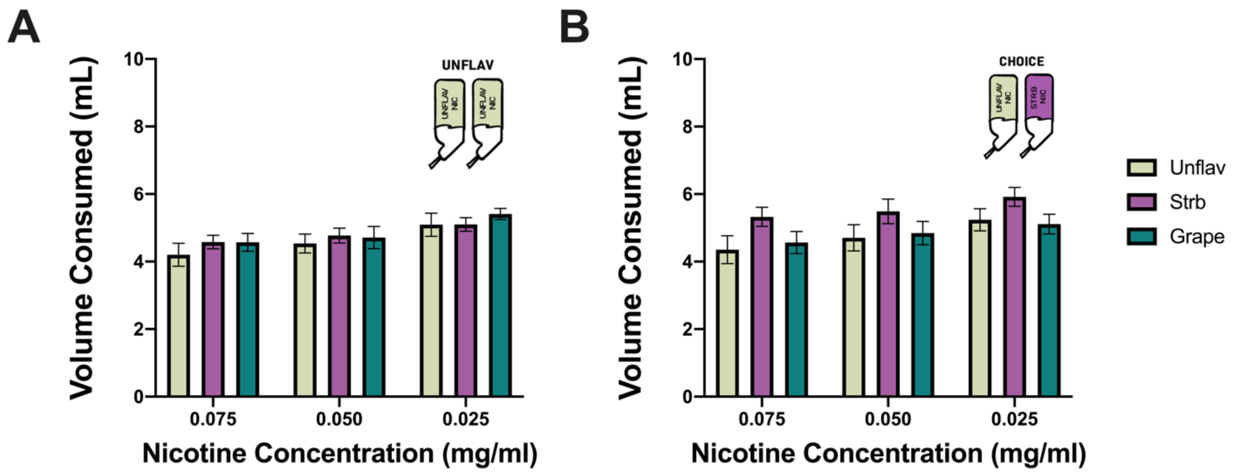


**Mice in all fading flavor groups drink equivalent volumes of fluid across all concentrations of nicotine.** Bar graphs represent summary data (mean +/- SEM). A) Mice that were in the UNFLAV group drink statistically equivalent volumes of fluid, regardless of fading flavor assignment, throughout the "Fading and Flavor Reintroduction" experiment. There was a significant effect of nicotine concentration, but no effect of fading flavor or an interaction (Two-way ANOVA, $F_{\text{fading flavor}}(2,32) = 0.478$, p=0.624, $F_{\text{nicotine conc.}}(1.227, 39.26) = 11.52$, ***p=0.0008, $F_{\text{interaction}}(4, 64) = 0.352$, p=0.842). B) Mice that were in the CHOICE group (i.e. had access to strawberry nicotine in adolescence) also drink equivalent volumes of fluid, regardless of fading flavor assignment, throughout the "Fading and Flavor Reintroduction" experiment. (Two-way ANOVA, $F_{\text{fading flavor}}(2,32) = 1.893$, p=0.167, $F_{\text{nicotine conc.}}(1.610, 51.53) = 18.44$, ****p<0.0001, $F_{\text{interaction}}(4, 64) = 0.561$, p=0.692). (UNFLAV in adolescence, 'No Flavor': n=12, 'Strawberry': n= 11, 'Grape': n=12; CHOICE in adolescence, 'No Flavor': n=10, 'Strawberry': n= 11, 'Grape': n=12)
